# Mendelian randomization suggests a bidirectional, causal relationship between physical inactivity and obesity

**DOI:** 10.1101/2021.06.16.448665

**Authors:** Germán D. Carrasquilla, Mario García-Ureña, Tove Fall, Thorkild I.A. Sørensen, Tuomas O. Kilpeläinen

**Affiliations:** Novo Nordisk Foundation Center for Basic Metabolic Research, Faculty of Health and Medical Sciences, University of Copenhagen, Copenhagen, Denmark; Molecular Epidemiology, Department of Medical Sciences, and Science for Life Laboratory, Uppsala University, Uppsala, Sweden; Department of Public Health, Section of Epidemiology, Faculty of Health and Medical Sciences, University of Copenhagen, Copenhagen, Denmark

**Author notes:** Corresponding author contact information: Name (Germán Darío Carrasquilla), work-phone, and postal mail address (Blegdamsvej 3B, Mærsk Tårnet, 8. sal, 2200 København N, Maersk Tower).

## Abstract

Physical inactivity is associated with excess weight gain in observational studies. However, some longitudinal studies indicate reverse causality where weight gain leads to physical inactivity. As observational studies suffer from reverse causality, it is challenging to assess the true causal directions. Here, we assess the bidirectional causality between physical inactivity and obesity by bidirectional Mendelian randomization analysis. We used results from genome-wide association studies for accelerometer-based physical activity and sedentary time in 91,105 individuals and for body mass index (BMI) in 806,834 individuals. We implemented Mendelian randomization using CAUSE method that accounts for pleiotropy and sample overlap using full genome-wide data. We also applied inverse variance-weighted, MR-Egger, weighted median, and weighted mode methods using genome-wide significant variants only. We found evidence of bidirectional causality between sedentary time and BMI: longer sedentary time was causally associated with higher BMI [beta (95%CI) from CAUSE method: 0.11 (0.02, 0.2), P=0.02], and higher BMI was causally associated with longer sedentary time (0.13 (0.08, 0.17), P=6.3.×10^-4^). Our analyses suggest that higher moderate and vigorous physical activity are causally associated with lower BMI (moderate: -0.18 (-0.3,-0.05), P=0.006; vigorous: -0.16 (-0.24,-0.08), P=3.8×10^-4^), but indicate that the association between higher BMI and lower levels of physical activity is due to horizontal pleiotropy. The bidirectional, causal relationship between sedentary time and BMI suggests that decreasing sedentary time is beneficial for weight management, but also that targeting obesity may lead to additional health benefits by reducing sedentary time.

## Background

Obesity and physical inactivity are major risk factors for a number of chronic diseases, such as type 2 diabetes, cardiovascular diseases and several types of cancer. Today’s epidemic of obesity and sedentary lifestyle are thus a major burden on public health systems worldwide (Collaborators, 2020).

Many observational studies suggest that physical inactivity is associated with a higher risk of obesity (Lee et al., 2010, Du et al., 2013, Silva et al., 2019, Myers et al., 2017). However, other studies have indicated a reverse effect, where obesity leads to physical inactivity (Petersen et al., 2004, Mortensen et al., 2006, Bak et al., 2004, Barone Gibbs et al., 2020, Ekelund et al., 2008, Myers et al., 2017). Furthermore, randomized clinical trials of physical activity interventions have indicated that the causal effects of physical activity on body weight are modest (Church et al., 2009, Rosenkilde et al., 2012, Golubic et al., 2015) compared to the strong inverse relationship between physical activity and body weight observed in cross-sectional epidemiological studies. This suggests that the observational results may be affected by bias, such as reverse causality or confounding by other lifestyle or environmental factors (Schnurr et al., 2021). To date, the causal relationships between physical inactivity and obesity remain unclear and warrant further investigation.

Mendelian randomization is a powerful method to minimize the influence of reverse causality and confounding on causal estimates derived from observational data. Since genotypes are randomly allocated at conception, genetic alleles associated with physical activity, sedentary behavior, and body mass index (BMI) can be used to assign individuals according to higher or lower mean levels of these exposures in a randomized manner.

Here, we aimed to assess the causality between physical inactivity and BMI by applying bidirectional Mendelian randomization analyses on summary results of accelerometer-based physical activity and sedentary time for 91,105 adults and of BMI for 806,834 adults.

## Methods

### Data sources and populations

We used summary results from the largest published genome-wide association studies (GWAS) of objectively assessed physical activity, sedentary behavior, and BMI in individuals of European ancestry. The physical activity GWAS included up to 91,105 individuals for accelerometer-based vigorous physical activity, moderate physical activity, or sedentary time from the UK Biobank (Klimentidis et al., 2018, Doherty et al., 2018). In these studies, accelerometer was worn continuously for at least 72 hours and up to 7 days. Vigorous physical activity was defined as the fraction of accelerations >425 milli-gravities [15], and moderate physical activity was predicted using a machine-learning method for moderate intensity activity time (Doherty et al., 2018). Sedentary time was defined as the time spent in activities with metabolic equivalent of task (MET) ≤ 1.5 during sitting, lying, or in reclining posture, except for driving and certain non-desk work instances where MET ≤ 2.5 was applied (Doherty et al., 2018). For BMI, we utilized GWAS results from a meta-analysis of the Genetic Investigation of Anthropometric Traits (GIANT Consortium) and the UK Biobank data, including altogether 806,834 individuals of European ancestry (Pulit et al., 2019). For Mendelian randomization analyses using the inverse variance-weighted (IVW), weighted median, weighted mode, and MR-Egger regression methods, we used only the GIANT Consortium BMI meta-analysis data of 339,224 individuals without the UK Biobank data to avoid sample overlap between the exposure and outcome traits as these methods are sensitive to bias from overlapping samples (Locke et al., 2015).

### Mendelian randomization using full genome-wide summary results for the exposure trait

Only few genetic loci have been found to be associated with accelerometer-based moderate physical activity (n=2), vigorous physical activity (n=1) or sedentary time (n=4) at genome-wide significance (P<5×10^-8^) (Klimentidis et al., 2018, Doherty et al., 2018), and the loci thus provide a limited power to study causal associations with BMI using Mendelian randomization. The recently published Causal Analysis Using Summary Effect Estimates (CAUSE) Mendelian randomization method (Morrison et al., 2020) improves statistical power in such cases, by utilizing full genome-wide summary results instead of genome-wide significant loci only. Furthermore, the CAUSE method is able to correct for sample overlap between the exposure and the outcome trait, which allows using the largest sample sizes available for both traits. CAUSE has also been found to be less prone to identify false positive associations compared to other commonly used Mendelian randomization methods (Burgess et al., 2019, Morrison et al., 2020).

The CAUSE method calculates the posterior probabilities of the causal effect and the shared (non-causal) effect, where the causal effect reflects the effect of the variants on the outcome trait through the exposure and the shared effect reflects correlated horizontal pleiotropy (Figure 1), i.e. the effect of the variants on the outcome through confounders. The distinction between a causal effect and correlated horizontal pleiotropy follows the assumption that a causal effect leads to non-zero genetic correlation between the exposure and the outcome where the correlation is driven by all variants associated with the exposure. If only a subset of variants contributes to the genetic correlation between the exposure and the outcome, it is considered the result of correlated horizontal pleiotropy. The CAUSE method also provides an estimate of the proportion of variants that are likely to show correlated horizontal pleiotropy, the q value.

**Figure 1.**
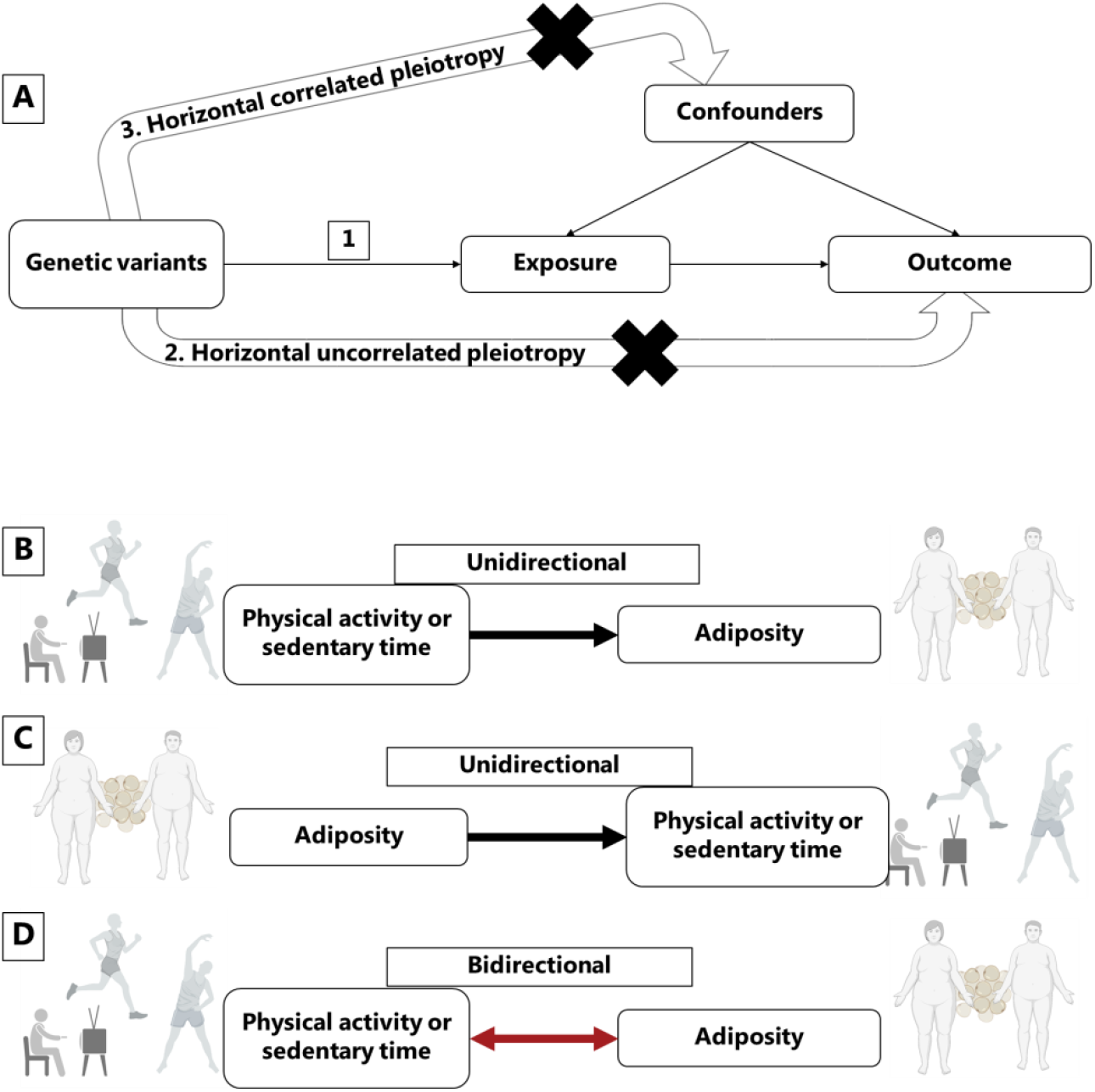
Mendelian randomization assumptions and directional associations between physical activity, sedentary time and adiposity. Panel A shows Mendelian randomization assumptions when estimating the causal association between a given exposure and outcome: 1) instrumental variants are associated with the exposure, 2) the instruments do not cause the outcome independently from the exposure (horizontal uncorrelated pleiotropy), and 3) the effects of the exposure on the outcome are not influenced by any confounders (horizontal correlated pleiotropy). Panel B indicates the one-way directional association between activities and adiposity, whereas panel C indicate unidirectional directions between adiposity and activities, here referred as reverse causation. Panel D indicates two-way directional associations where activities have an impact on adiposity, but at the same time adiposity impact levels of activity in a bidirectional manner. Figure icons were created with BioRender.com.

We used the CAUSE settings and procedures originally recommended by the authors (Morrison et al., 2020), with the exception of q priors that were set to fit the strictest model possible (q_alpha=1 and q_beta=2) in order to avoid false positive findings. A thorough explanation of the steps used to perform CAUSE analysis is included in the in the Supplementary text (Additional file 1).

### Mendelian randomization using genome-wide significant loci for the exposure trait

In addition to the CAUSE method that implements Mendelian randomization analyses using full genome-wide summary results for the exposure trait, we implemented four commonly used Mendelian randomization methods that utilize genome-wide significant loci only: the IVW, MR-Egger, weighted median and weighted mode methods (Additional file 1). We performed sensitivity analyses using Steiger filtering to remove variants that showed stronger association with the outcome than the exposure trait and that were thus not considered suitable as instruments for the exposure trait. To create the genetic instrument for the exposure trait, we only included the lead variants that showed genome-wide significant associations with the trait (P < 5×10^-8^) and with a pairwise linkage disequilibrium (LD) r^2^ < 0.001 with their neighboring variants, in a window of 10000kb. Variants that were not available in the outcome trait GWAS were substituted by their LD proxies (r^2^ > 0.8). Palindromic variants (A/T, G/C) were excluded. If less than three genetic variants were identified with these parameters, we used a less stringent p-value threshold of P < 5×10^-7^ to identify enough genetic instruments. In order to assess the strength of the genetic instrument, we obtained *F*-statistics for each trait. The analyses were performed using the TwoSampleMR package in R and are described in detail in the Additional file 1 (Hemani et al., 2018).

We estimated heterogeneity across the causal estimates of the SNPs using the Meta R package (Schwarzer et al., 2015). The causal estimates were considered heterogeneous if the P value for Cochran’s Q test was significantly different from zero (P<0.05) and I^2^ was above 0.25. We assessed bias introduced by horizontal pleiotropy by implementing the Egger’s intercept test using the TwoSampleMR package in R (Hemani et al., 2018). An Egger’s intercept that deviated significantly from zero (P<0.05) was considered as evidence of horizontal pleiotropy. We used the Rucker framework (Bowden et al., 2018) to assess whether Egger regression that accounts for horizontal pleiotropy but limits statistical power should be applied instead of the standard IVW model. To visually assess heterogeneity and horizontal pleiotropy, we observed forest plots and funnel plots (Supplementary Figure S1). To detect individual pleiotropic variants that might bias the results, we applied the RadialMR package in R using an iterative Cochran’s Q method and setting a strict outlier P value threshold of <0.05 (Bowden et al., 2018). The iterative Cochran’s Q, either IVW’s Q or Egger’s Q’ was chosen depending on Rucker framework results. After removing outlier variants detected with RadialMR, we re-run the Mendelian randomization and sensitivity test and plots, to make sure that the variants introducing horizontal pleiotropy (Figure 1) had been removed. The analysis plan for this study is described in the in the Supplementary text (Additional file 2).

The CAUSE method’s median posterior probability of the causal effect cannot be easily transformed to absolute units. To convert the causal estimates to absolute units, we calculated a causal effect with weighted median method using independent variants identified in CAUSE that were not removed by the outlier extraction protocol described above, in order to mimic CAUSE control for correlated and uncorrelated pleiotropy (Additional file 1).

## Results

We use the Mendelian randomization CAUSE method to take advantage of the full genome-wide summary results.(Morrison et al., 2020) We found evidence of causal association between higher vigorous and moderate physical activity and lower BMI (P=3.8×10^-4^ and P=0.006, respectively), and between more sedentary time and higher BMI (P=0.02) (Table 1, Figure 2, and Supplementary Table S1 and S2). The median shared effect ranged from -0.01 to 0, which reflects the effect induced by correlated horizontal pleiotropy, was zero for all trait pairs, indicating that there was no bias induced by horizontal pleiotropy. The low q values (q=0.19-0.20), which reflect the proportion of variants that show correlated horizontal pleiotropy, also suggested that horizontal pleiotropy was limited. In absolute units, we approximate that each one hour daily increase in moderate physical activity or decrease in sedentary time was causally associated with 0.27 kg/m^2^ (∼0.8 kg) or 0.14 kg/m^2^ (∼0.4 kg) lower BMI, respectively (Supplementary Table S5).

**Table 1.** Results for Mendelian randomization analyses using the CAUSE method.

|  |  |  |  |
| --- | --- | --- | --- |
| Causal model better fit for the data |  |  |  |
| Direction | Median causal effect (95% CI) | Median q (CI) | P causal vs sharing |
| Vigorous PA → BMI | -0.16 (-0.24, -0.08) | 0.19 (0, 0.86) | 3.8x10<sup>-04</sup> |
| Moderate PA → BMI | -0.18 (-0.3, -0.05) | 0.20 (0.01, 0.86) | 0.006 |
| Sedentary time → BMI | 0.11 (0.02, 0.20) | 0.19 (0, 0.86) | 0.02 |
| BMI → Sedentary time | 0.13 (0.08, 0.17) | 0.18 (0, 0.85) | 6.3x10<sup>-4</sup> |
| Sharing model better fit for the data |  |  |  |
| Direction | Median shared effect (CI) | Median q (CI) | P causal vs sharing |
| BMI → Vigorous PA | -0.18 (-0.3, -0.05) | 0.94 (0.77, 0.98) | 0.35 |
| BMI → Moderate PA | -0.11 (-0.19, -0.03) | 0.77 (0.55, 0.95) | 0.31 |
| The results display the data according to the goodness-of-fit for the causal or the sharing model. The median q value indicates the proportion of variants with correlated pleiotropy. CI, confidence interval; PA, physical activity; BMI, body mass index; P, P-value. |  |  |  |

**Figure 2.**
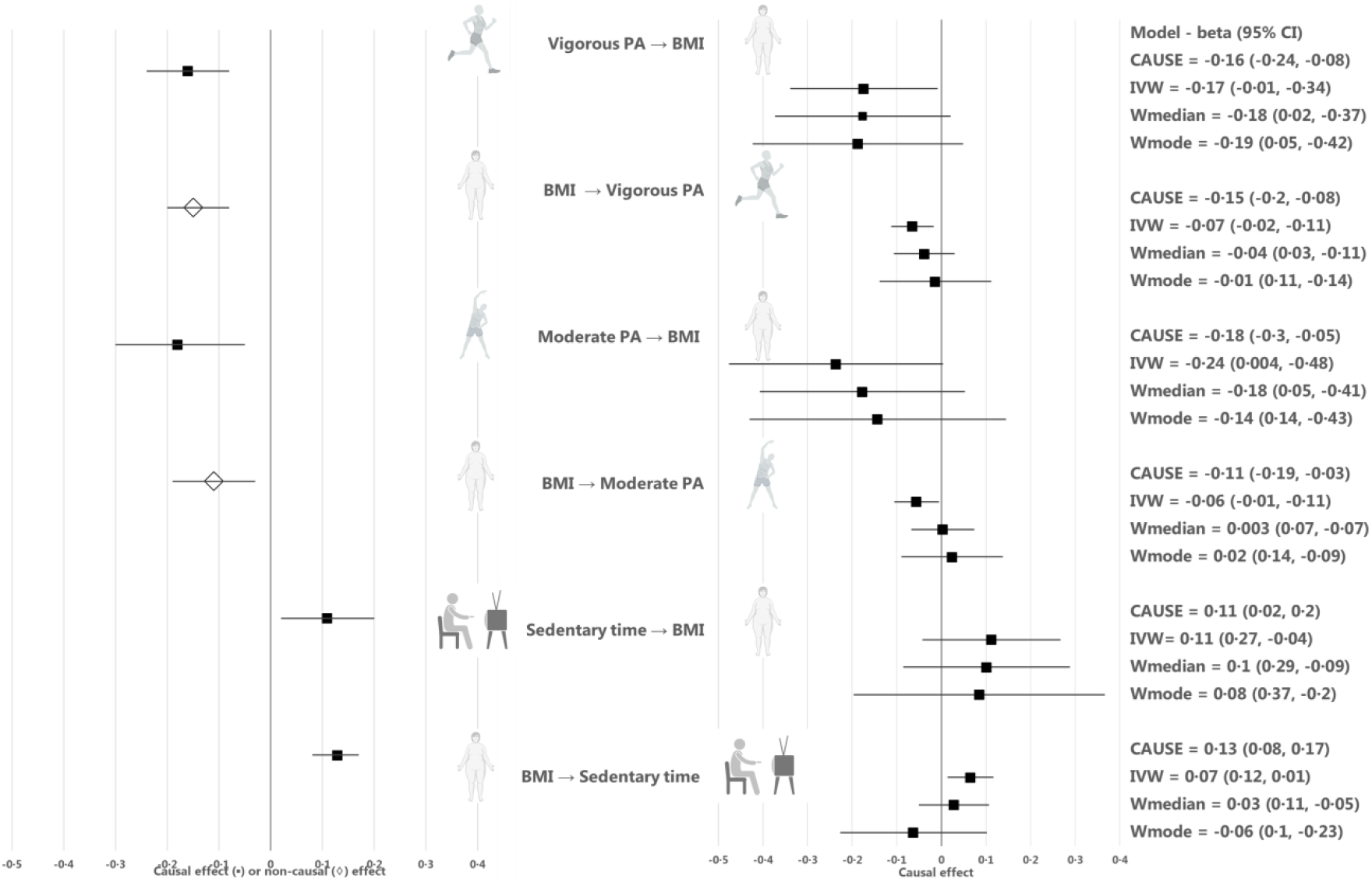
Causal estimates for Mendelian randomization analyses using the CAUSE, inverse-variance-weighted (IVW), MR Egger regression, weighted median, weighted mode methods. Median causal estimates for Mendelian randomization analyses using the CAUSE method (left panel) and mean causal estimates from the inverse variance weighted (IVW), weighted median (Wmedian) and weighted mode (Wmode) methods, after outlier removal and accounting for horizontal pleiotropy. A diamond (◊) in the estimate for CAUSE indicates that the sharing model fit the data better than the causal model, i.e. that the association between the traits was more likely to be explained by horizontal correlated pleiotropy than causality. PA, physical activity; BMI, body mass index. Figure icons were created with BioRender.com

In the reverse direction, we found no evidence of a causal effect of BMI on vigorous physical activity (P=0.35) or moderate physical activity (P=0.31) using CAUSE (Table 1, Figure 2, and and Supplementary Table S1 and S2). However, we found evidence of causal effects of BMI on more sedentary time (P=6.3×10^-4^), indicating bidirectional causality between the traits. The median shared effect in the causal association between BMI and sedentary time was zero and the q value was 0.18, suggesting that the causal association between sedentary time and BMI was unlikely to be biased by horizontal pleiotropy. In absolute units, we approximate that each kg/m^2^ (∼3 kg) increase in BMI was causally associated with a 3.5 minute increase in sedentary time per day (Supplementary Table S5).

We also estimated the causal effects of moderate physical activity, vigorous physical activity and sedentary time on BMI with four commonly used Mendelian randomization methods, including IVW, Egger, weighted median and weighted mode methods. Due to the low number of independent, genome-wide significant loci for vigorous physical activity, moderate physical activity and sedentary time that were present in the GWAS results for BMI, we used a less stringent threshold of P<5×10^-7^ to identify genetic instruments for these traits, resulting in 5, 3 and 5 independent loci, respectively. The directions of causal estimates were consistent with the findings from CAUSE, but the evidence for causality was weaker (Table 2, Figure 2, Additional file 1, and Supplementary Table S3). To estimate the causal effect of BMI on moderate physical activity, vigorous physical activity and sedentary time, we used genome-wide significant BMI loci (P<5×10^-8^) as instruments (n=57, n=55 and n=57, respectively). Again, the directions of causal estimates were consistent with the CAUSE results, but the associations were weaker (Table 2, Figure 2, Additional file 1, and Supplementary Table S3).

**Table 2.** Mendelian randomization results for inverse variance weighted, weighted median, weighted mode, and MR-Egger 219 methods.

| Table 2. Mendelian randomization results for inverse variance weighted, weighted median, weighted mode, and MR-Egger, methods |  |  |  |  |  |  |  |  |  |
| --- | --- | --- | --- | --- | --- | --- | --- | --- | --- |
| Direction | Vigorous physical activity → BMI |  |  | Moderate physical activity → BMI |  |  | Sedentary time → BMI |  |  |
| MR method | beta | SE | p-value | beta | SE | p-value | beta | SE | p-value |
| IVW | -0.17 | 0.08 | 0.04 | -0.24 | 0.12 | 0.05 | 0.11 | 0.08 | 0.16 |
| Weighted median | -0.18 | 0.10 | 0.08 | -0.18 | 0.12 | 0.13 | 0.10 | 0.10 | 0.29 |
| Weighted mode | -0.19 | 0.12 | 0.19 | -0.14 | 0.15 | 0.43 | 0.08 | 0.14 | 0.59 |
| MR-Egger | 1.33 | 2.21 | 0.59 | 0.12 | 0.45 | 0.84 | 0.49 | 0.36 | 0.26 |
| Direction | BMI → Vigorous physical activity |  |  | BMI → Moderate physical activity |  |  | BMI → Sedentary time |  |  |
| IVW | -0.07 | 0.02 | 0.01 | -0.06 | 0.03 | 0.03 | 0.07 | 0.03 | 0.01 |
| Weighted mode | -0.01 | 0.06 | 0.83 | 0.02 | 0.06 | 0.68 | -0.06 | 0.08 | 0.46 |
| Weighted median | -0.04 | 0.03 | 0.27 | 0.003 | 0.04 | 0.93 | 0.03 | 0.04 | 0.48 |
| MR-Egger | 0.13 | 0.07 | 0.06 | 0.16 | 0.07 | 0.04 | -0.02 | 0.08 | 0.76 |
| BMI, Body mass index (BMI); SE, standard error; IVW, inverse variance weighted; CI, 95% confidence interval. |  |  |  |  |  |  |  |  |  |

## Discussion

The present Mendelian randomization analyses suggest a bidirectional causal relationship between higher sedentary time and higher BMI, implying that decreasing sedentary time is beneficial for weight management, but also that reducing obesity may lead to additional health benefits by reducing sedentary time. The analyses also suggest there is a causal association between higher levels of physical activity and lower BMI, supporting the view that preventive programs for increasing physical activity and decreasing sedentary time are beneficial for weight management.

Our results are in accordance with a previous observational study that aimed to assess the bidirectional relationship between physical activity and weight change during a 10-year period.(Barone Gibbs et al., 2020) Examining associations between accelerometer-based activity measures and weight change in 866 men and women, the study suggested a bidirectional relationship where higher sedentary behavior at baseline increased 10-year weight gain and higher baseline weight was associated with an unfavorable 10-year change in sedentary time. Our results are also in accordance with randomized clinical trials of physical activity interventions which generally suggest that increasing physical activity leads to a moderate loss of body weight in overweight or obese participants (Church et al., 2009, Rosenkilde et al., 2012, Golubic et al., 2015, Kim et al., 2019, Biddle et al., 2015). Based on the causal effect size, we estimated that one hour daily increase in moderate physical activity leads to ∼0.8 kg decrease in body weight (Additional file 1 and Supplementary Table S5). However, it is important to note that the causal estimates from Mendelian randomization are not fully comparable to those from randomized clinical trials, because they represent lifelong effects rather than effects lasting a defined length of an intervention, and furthermore, physical activity interventions may operate on body weight through other pathways than those affected by the genotypes. The causal association between higher adiposity and physical inactivity has not been to date assessed in randomized clinical trials, likely due to the ethical and practical limitations of performing such a study.

In a previous Mendelian randomization analysis of adult populations, evidence for a causal, bidirectional relationship between overall activity levels and higher BMI was observed using the maximum likelihood method, but the results showed evidence of horizontal pleiotropy that could not be fully accounted for and the role of activity intensity level remained unclear (Doherty et al., 2018). Here, using a method that takes advantage of full genome-wide summary results and corrects for sample overlap between the exposure and the outcome traits to maximize statistical power and correct for pleiotropy, we showed that the causal bidirectional relationship is particularly evident for the relationship between sedentary time and obesity. Our results may also be compared with two independent one-sample Mendelian randomization studies performed in children (Richmond et al., 2014, Schnurr et al., 2018). The first study, including 4,296 children at 11 years of age from the United Kingdom, indicated a causal association between higher BMI and lower accelerometer-based moderate and moderate-to-vigorous physical activity and more sedentary time (Richmond et al., 2014). The second study, including 679 children at age 3-8 years from Denmark, also indicated that higher BMI is causally associated with higher accelerometer-based sedentary time, but did not find a causal association with moderate or moderate-to-vigorous physical activity (Schnurr et al., 2018). Consistent with the latter study of children, our results indicate a causal effect of BMI on sedentary behavior, but not on physical activity, in adults. The differences between studies could be due to different applied methods, or methodological limitations, such as weak instrument bias when smaller sample sizes are used, which may lead to estimated causal effects towards the observational association. One could also expect differences between children and adults given the distinct patterns by which they engage in physical activity. For example, while physical activity in adults consists of commuting, occupational and structured leisure-time activities, children primarily engage in spontaneous, play-oriented activities. Higher BMI leads to higher perceived exertion during physical activity (Groslambert and Mahon, 2006), which could reduce the natural inclination of children to engage in play-oriented activities, whereas adults exert more conscious control over their daily activities.

The strengths of the present studies include the use of genome-wide summary results for objectively measured physical activity and sedentary time, which avoided misreporting bias evident for self-reported measures, as well as the use of newly developed Mendelian randomization method that utilizes full genome-wide summary results to improve statistical power, correct for sample overlap, and assess horizontal pleiotropy, successfully applied in recent Mendelian randomization studies (Jager et al., 2020, Mitchell et al., 2020). The limitations are that we cannot exclude other sources of bias in the measurement of physical activity and sedentary time that could influence the observed causal estimates, including the observer effect and the limited 7-day period of the measurement, which may not be representative of long-term activity habits. Furthermore, even if we used the largest available data on objectively measured physical activity, the statistical power was limited, as very few genome-wide significant loci have thus far been identified.

## Conclusion

The present Mendelian randomization analyses indicate a bidirectional causal relationship between higher sedentary time and higher BMI. Thus, increasing sedentary time is likely to be beneficial for weight management, but reducing obesity may also lead to additional health benefits by reducing sedentary time. Our analyses also suggest that there is a causal association between higher levels of physical activity and lower BMI, supporting the view that lifelong preventive programs for increasing physical activity and decreasing sedentary time are beneficial for weight management.

## Acknowledgments

This project has received funding from the European Union’s Horizon 2020 research and innovation programme under the Marie Sklodowska-Curie grant agreement No 846502. Novo Nordisk Foundation Center for Basic Metabolic Research is an independent research center at the University of Copenhagen partially funded by an unrestricted donation from the Novo Nordisk Foundation (NNF18CC0034900). Germán D. Carrasquilla was supported by a grant from the Danish Diabetes Academy that is funded by the Novo Nordisk Foundation (NNF17SA0031406). Tuomas O. Kilpeläinen was supported by a grant from the Novo Nordisk Foundation (NNF17OC0026848). The funding source had no role in study design, data collection, data analysis, data interpretation, or writing of the report. All authors had full access to all of the data in the study and had final responsibility for the decision to submit for publication.

## Additional file 1. Mendelian randomization using the CAUSE, IVW, Egger, weighted median, and weighted mode methods

### Mendelian randomization using the CAUSE method

The CAUSE method performs Mendelian randomization analyses following six different steps, as described in the CAUSE online tutorial [1] and the original publication [2]. The steps are the following:

1. Installing the followiing versions of these three packages were used: CAUSE v1.0.0, mixsqp v.0.1-97 and ashr v.2.2-32.
2. Filtering data by including variants with imputation quality score INFO > 0.7 and minimum allele frequency (MAF) > 0.01.
3. Excluding variants from the Major Histocompatibility Complex (MHC) present in chromosome 6 between the base pairs 26M and 34M in build 37.
4. Merging the exposure and outcome GWAS summary statistic level data. Gwas_merge function from CAUSE package was used to identify the variants present in exposure and outcome summary statistics data and to align exposure and the outcome effect sizes to the same allele
5. Calculating nuisance parameters to correct for sample overlap between exposure and outcome GWAS.
6. Using HapMap3 reference panel to select SNPs with LD r^2^ < 0.1.
7. Setting the priors for the three model parameters – causal effect, shared effect and q – and calculating their posterior probabilities. For the causal effect and the shared effect, their priors are set automatically to 0, while for q the software allows the user to set the thresholds for the priors. In this case, q priors are set to qalpha = 1 and qbeta = 2.
8. Calculating two models to fit the posterior probabilities: the sharing model, where the causal effect is set to 0, and the causal model, where the posterior probability for the causal effect is calculated.
9. Comparing the two models, sharing and causal, against the null and against each other with expected log pointwise posterior density (ELPD) method to identify which model is the most fitting for the data.

### Mendelian randomization using the IVW, Egger, weighted median, and weighted mode methods

We used the following parameters to interpret the findings:

1. Since the version of TwoSampleMR used in the analysis v0.5.4 removes duplicates by excluding the second instance when introducing data locally, only the variant that presented the same alleles as in the outcome data and with the lowest p-value were kept. In none of the combinations of traits, variants in the MHC were found.
2. To clump variants, the function ld_clump_local from the package ieugwasr v0.1.5 https://github.com/MRCIEU/ieugwasr) was used using the updated European 1000 Genomes reference panel available in https://github.com/mrcieu/gwasglue.
3. Only variants with MAF > 0.01 and INFO > 0.7 were included in the analysis.
4. Only variants that passed the Steiger filtering using the function steiger_filtering from the package TwoSampleMR were used.
5. The NOME and InSIDE assumptions were checked calculating the mean F-statistic and the variation of the I^2^ statistic, respectively. The later can be calculated with the Isq function from TwoSampleMR package.
6. The causal estimates were considered heterogeneous if the P value for Cochran’s Q test was <0.05 and I^2^ was >0.25. Both estimates were calculated using the meta package.
7. An Egger’s intercept P value <0.05 was considered as evidence of horizontal pleiotropy.
8. To assess whether MR-Egger regression should be applied instead of the standard IVW model, we used the Rucker framework test.
9. To detect individual pleiotropic variants, we used RadialMR’s iiterative Cochran’s Q method following a P value threshold <0.05. RadialMR presents two functions: ivw_radial or egger_radial, depending on the Cochran’s Q, either IVW’s Q or Egger’s Q’, used. The function used was chosen depending on Rucker framework test’s result. If Rucker framework presented contradictory results, an iterative version of the Rucker framework (i.e. rucker_jackknife from TwoSampleMR package) was used to assess whether IVW was still chosen as the main model.

To visualize the effects of heterogeneity and horizontal pleiotropy on the results, we generated leave-one-out forest plots and funnel plots. After removing outlier variants detected with RadialMR, we re-run the Mendelian randomization methods and sensitivity tests (eTable 3 and 4) and re-generated the plots (eFigure 1), to make sure that the variants introducing horizontal pleiotropy had been removed. Since the meanThe chances of weak instrument was low given an *F*-statistics ranged between 28,6 to 61,79 (F statistic > 10) indicate that results are based on valid instrumental variables, meaning that weak instrument bias may not be an issue. Furthermore, we tested whether the amount of pleiotropy was independent of instrument strength by calculating a variation of the I^2^ (Bowden et al., 2019) that was above 0..90 in all cases. Below we describe and interpret the findings for each combination of traits.

### Vigorous physical activity → BMI

All MR methods except Egger showed negative causal effects, from which only IVW is significant (P<0.05) (eTable 3). No outlier extraction was performed since Q’s test was not significant (P=0.80) and low I^2^ (0.0%) showed no evidence of heterogeneity. Egger’s intercept was not significant (P=0.30) and Rucker test indicated that IVW model is more fitting for the data (eTable 3). Consistent with the sensitivity tests, leave-one-out forest plot (eFigure 1, panel a) showed that the IVW causal effect did not strongly change after the removal of any of the five independent variants used to calculate the causal effects. Funnel plot (eFigure 1, panel a) showed no signs of asymmetry, which is consistent with Egger’s intercept result. Thus, these findings indicate that the causal effects are not biased and they reflect weak evidence of causality between an increase of vigorous physical activity and a decrease of BMI. While IVW negative causal effect is significantly different from 0, the evidence for causality is considered weak due to 1) IVW’s p-value is still close to nominal threshold (P=0.04), 2) the other methods with negative causal effects are not significantly different from 0 (P>0.05) and 3) Egger’s regression presents a positive causal effect. These weak results are most probably due to the low number of variants used (eTable 3).

### Moderate physical activity → BMI

All MR methods except Egger showed negative effects directions, with only IVW’s being significant (eTable 3). No outlier extraction was performed since Q’s test was not significant (P=0.30) and I^2^ was below 25% (18.9%). Egger’s intercept was not significantly different from 0 (P=0.16) indicating that no pleiotropic variants were found among the four independent variants used to calculate the causal effects. Rucker tests indicated that IVW was a better fit for the model than Egger, in line with Egger’s intercept results (eTable 3). The leave-one-out forest sensitivity plot (eFigure 1, panel c) showed that no variant presented heterogenic causal effect since the IVW causal effect did not strongly diverge after removing any of the variants. The funnel sensitivity plot (eFigure 1, panel c) showed no asymmetry. All in all, the findings indicate that the causal effects are not biased, but that they suggest causality between increased moderate physical activity and decreased BMI. The reason behind this is that 1) IVW’s negative causal effect present a p-value close to the nominal threshold (P=0.02), 2) the other methods present negative causal effect that are not significantly different form 0 (P>0.05) and 3) Egger’s regression presents a positive causal effect, though it is not significantly different from 0 (eTable 3).

### Sedentary time → BMI

Nine independent variants were used as instruments, and two were removed after outlier extraction as indicated by RadialMR. All methods were not significant, and all reported a positive causal effect (P>0.05), except weighted mode (eTable 3). After outlier extraction, Cochran’s Q test was non-significant (P=0.15), but the I^2^ was 36.4%, indicating slight heterogeneity. Egger’s intercept was not significant (P=0.43) and Rucker test indicated that IVW method is a better fit for the data than Egger, implying that pleiotropic variants were not present (eTable 3). In accordance, the forest plot and funnel plot (eFigure 1, panel e) showed that the results are not strongly affected by horizontal pleiotropy. The forest plot indicated that the heterogeneity comes from variants with both, positive and negative causal effects, since the IVW causal effect becomes either more strongly negative or positive when the variants are removed one at a time. The funnel plot (eFigure 1, panel e) showed that the causal estimates are evenly distributed, implying no asymmetry and, hence no pleiotropy. The lack of strong variants with low standard errors is probably the reason behind heterogeneity of the data. To conclude, the findings indicate a non-significant positive causal effect between sedentary time and BMI.

### BMI → Vigorous physical activity

All MR methods except Egger presented negative causal estimates. The causal estimate from the IVW method reached statistical significance (P<0.01) (eTable 4). Non-significant P value from Cochran’s Q’ s test (p = 0.27) and low I^2^ of 9.6% indicated no heterogeneity. Egger intercept’s was significant (P=0.0017) and IVW’s Q and Egger’s Q were significantly different (Q-Q’ = 9.83, p= 0.0017), indicating that horizontal pleiotropy may still affect the causal estimates. Rucker framework selected IVW more fitting for the data than Egger (eTable 4). Forest and funnel plots (eFigure 1, panel b) indicated that some variants, both with positive and negative effects, may be introducing heterogeneity. The forest plot showed slight deviations from the mean causal effect when extreme variants were extracted one at a time. In the funnel plot, symmetry was disrupted by variants with positive causal effects and small standard errors. Considering both plots, we infer that heterogeneity and horizontal pleiotropy were introduced by specific variants with small standard errors. Leave-one-out plots indicated that two variants (rs13021737 and rs6567160) introduced horizontal pleiotropy, showing positive causal effects and having the smallest standard errors of all variants. To conclude, we found residual pleiotropy that RadialMR could not properly address and deem that the association between BMI and vigorous physical activity is unlikely to be causal.

### BMI → Moderate physical activity

The IVW method showed a negative causal effect, whereas Egger, weighted median and weighted mode showed a positive causal effect (eTable 4). Only the causal effect from Egger was significant (P=0.04). Non-significant P from Cochran’s Q test (P=0.91) and I^2^ of 0% indicated no presence of heterogeneity. Egger intercept’s was significant (P=0.002) and IVW’s Q and Egger’s Q were significantly different (Q-Q’ = 9.60, p= 1.95×10^-5^), indicating that horizontal pleiotropy may still affect the causal estimates. Rucker framework chose IVW as the most fitting method for the data. Forest and funnel plots (eFigure 1, panel d) indicated heterogeneity. The leave-one-out forest plots indicated deviation from the mean causal effect when variants with positive causal effects were excluded, and the funnel plots showed that asymmetry was introduced by variants with positive causal effects and small standard errors. These findings were in agreement with the findings from Rucker framework and Egger’s intercept test results. To conclude, we found evidence of horizontal pleiotropy that RadialMR could not properly remove, and where pleiotropy was introduced by variants with positive causal estimates. The presence of horizontal pleiotropy may explain the negative direction of the causal estimate in the IVW method, whereas other methods showed positive causal estimates (eTable 4). We deem that the association between BMI and moderate physical activity is unlikely to be causal.

### BMI → sedentary time

The IVW and weighted median methods showed positive causal effects, while Egger and weighted mode presented negative causal effects (eTable 4), with only IVW being significant (P= 0.02). The Cochran’s Q’ s test (P=0.4) and I^2^ of 2.2% indicated no presence of heterogeneity. Egger intercept test (P=0.20) and the non-significant difference between IVW’s Q and Egger’s Q (Q-Q’ = 1.62, p= 0.20) indicated no presence of horizontal pleiotropy and that the IVW method was a better fit for the data than Egger (eTable 4). Forest and funnel sensitivity plots (eFigure 1, panel f) indicated that there was no heterogeneity, except for one variant (rs6567160) with a negative effect. In the forest plot, upper and lower extreme variants induced deviations from the mean causal effect when extracted one at a time. To conclude, sensitivity tests indicate no horizontal pleiotropy and forest and funnel plots indicate the presence of heterogeneity for one variant, while RadialMR could not detect any outliers. Taken together, even though Rucker framework selected IVW as better fit for the data than Egger, the forest and funnel plots indicated heterogeneity, implying that there was residual horizontal pleiotropy. Therefore, we deem that there is no causal relationship between BMI and sedentary time, despite the significant P value in the IVW method.

### Conversion of the causal estimates to absolute units

To interpret the causal estimates from CAUSE, we applied the weighted median method on the independent instrumental variants selected by the CAUSE method and performed outlier extraction and sensitivity tests, following the steps described below. The results are reported in eTable 5.

1. We selected independent variants associated with each exposure identified with the CAUSE method.
2. We performed outlier extraction using RadialMR to control for uncorrelated pleiotropy.
3. We obtained weighted median’s causal estimates, to correct for correlated pleiotropy.
4. We approximated the causal estimates between outcome and exposure in absolute units. As the genome-wide summary results were reported in standard deviation units, we multiplied the causal estimates by the standard deviation of the corresponding non-transformed trait to derive the estimates in original trait units. The equations used in these calculations are described for each combination of traits in eTable 5.

---

**Figure S1.**
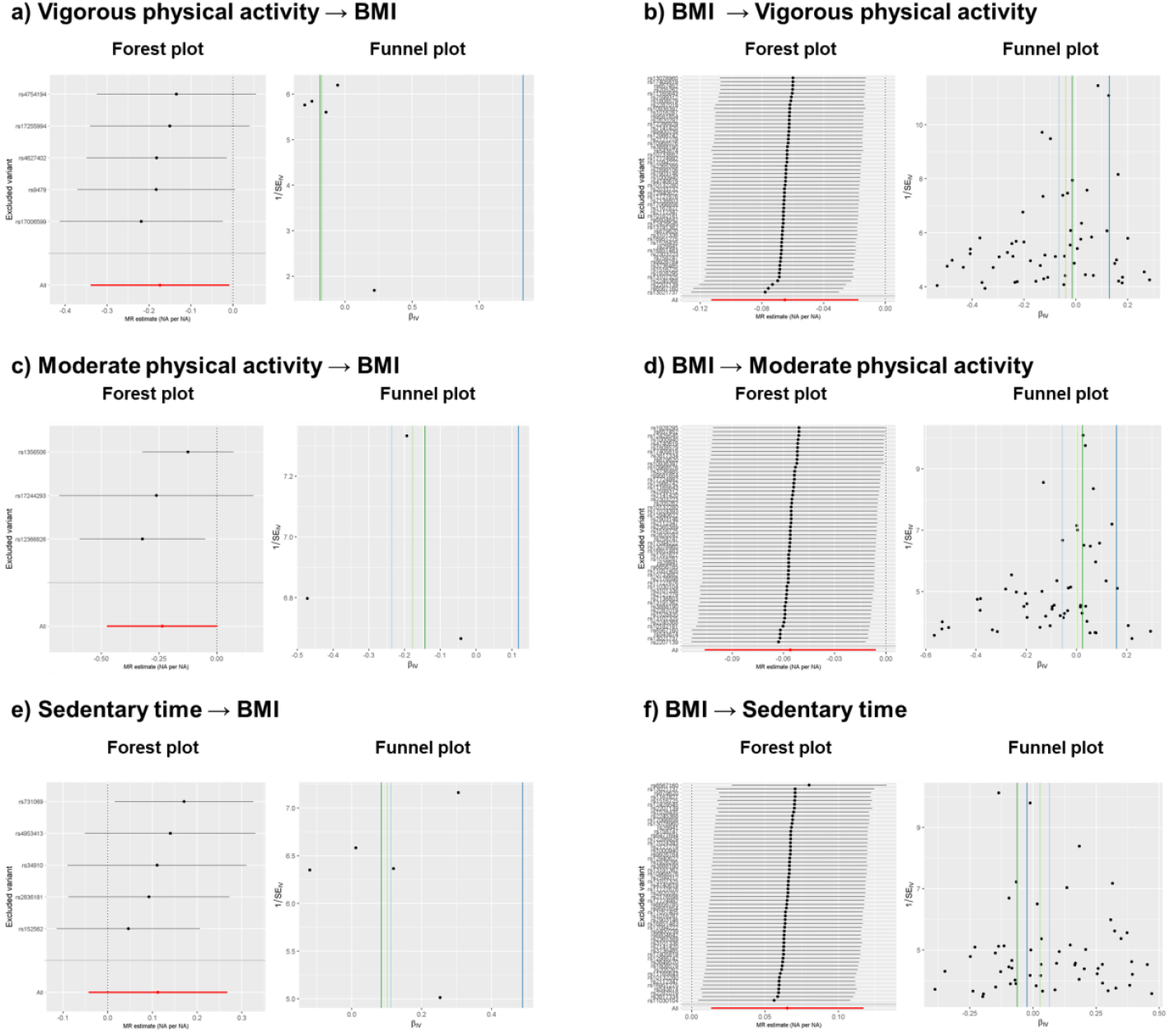
Leave-one-out forest and funnel sensitivity plots after outlier extraction.

**Table S1.** expected log pointwise posterior density (ELPD) results for each combination of traits using the CAUSE method.

| **Table S1: CAUSE expected log pointwise posterior density (ELPD) results for each combination of traits** |  |  |  |  |  |  |  |
| --- | --- | --- | --- | --- | --- | --- | --- |
| **Model 1** | **Model 2** | **Vigorous physical activity → BMI** |  |  | **BMI → Vigorous physical activity** |  |  |
|  |  | **Delta_ELPD** | **SE_delta_ELPD** | **z** | **Delta_ELPD** | **SE_delta_ELPD** | **z** |
| Null | Sharing | -17.47 | 4.24 | -4.12 | -251.88 | 21.54 | -11.69 |
| Null | Causal | -18.98 | 4.68 | -4.05 | -252.22 | 21.65 | -11.65 |
| Sharing | Causal | -1.51 | 0.45 | -3.36 | -0.35 | 0.90 | -0.38 |
|  |  | **Moderate physical activity → BMI** |  |  | **BMI → Moderate physical activity** |  |  |
|  |  | **Delta_ELPD** | **SE_delta_ELPD** | **z** | **Delta_ELPD** | **SE_delta_ELPD** | **z** |
| Null | Sharing | -8.50 | 2.86 | -2.97 | -119.39 | 15.15 | -7.88 |
| Null | Causal | -9.83 | 3.39 | -2.90 | -119.77 | 15.34 | -7.81 |
| Sharing | Causal | -1.34 | 0.53 | -2.52 | -0.38 | 0.77 | -0.49 |
|  |  | **Sedentary time → BMI** |  |  | **BMI → Sedentary time** |  |  |
|  |  | **Delta_ELPD** | **SE_delta_ELPD** | **z** | **Delta_ELPD** | **SE_delta_ELPD** | **z** |
| Null | Sharing | -6.51 | 2.57 | -2.53 | -137.84 | 157.46 | -8.75 |
| Null | Causal | -7.67 | 3.14 | -2.44 | -138.89 | 15.88 | -8.75 |
| Sharing | Causal | -1.16 | 0.57 | -2.05 | -1.06 | 0.33 | -3.22 |
| BMI, Body mass index; SE, standard error; NA, not applicable; CI, 95% confidence interval. |  |  |  |  |  |  |  |

**Table S2.**
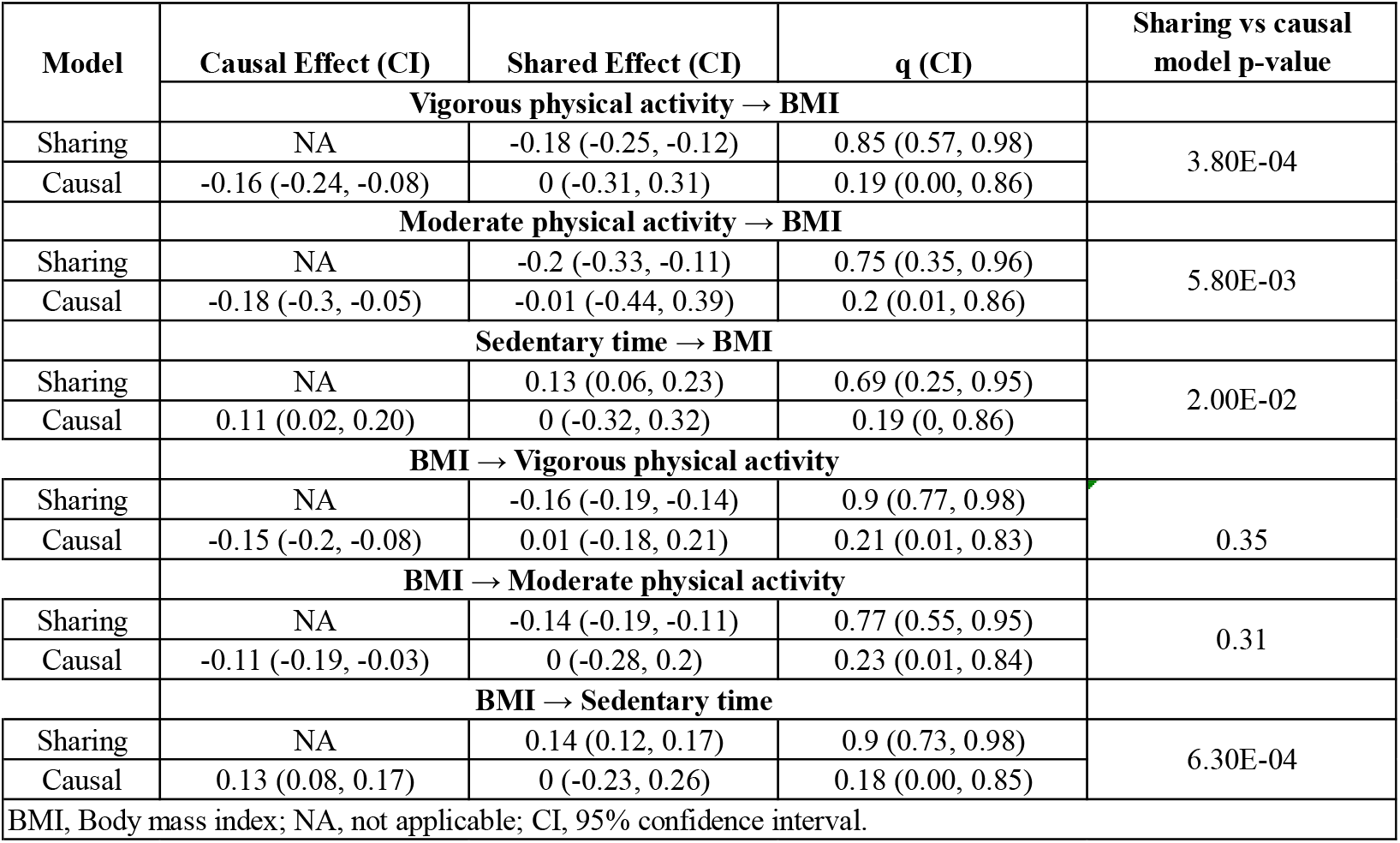
CAUSE posterior probabilities and q values for the causal effect and the shared effect.

**Table S3.** Mendelian randomization results using the IVW, Egger, weighted median and weighted mode methods of vigorous, moderate physical activity and sedentary on BMI

| Table S3. Mendelian randomization results using the IVW, Egger, weighted median and weighted mode methods of moderate, vigorous physical activity and sedentary time on BMI |  |  |  |  |  |  |  |  |  |  |  |  |
| --- | --- | --- | --- | --- | --- | --- | --- | --- | --- | --- | --- | --- |
| Before outlier extraction |  |  |  |  |  |  |  |  |  |  |  |  |
| MR method | Vigorous physical activity → BMI |  |  |  | Moderate physical activity → BMI |  |  |  | Sedentary time → BMI |  |  |  |
|  | SNP | beta | SE | p-value | SNP | beta | SE | p-value | SNP | beta | SE | p-value |
| Egger | 5 | 1.33 | 2.21 | 0.59 | 3 | 0.12 | 0.45 | 0.84 | 8 | 0.46 | 0.64 | 0.50 |
| Weighted median | 5 | -0.18 | 0.10 | 0.08 | 3 | -0.18 | 0.12 | 0.13 | 8 | 0.02 | 0.09 | 0.82 |
| IVW | 5 | -0.17 | 0.08 | 0.04 | 3 | -0.24 | 0.12 | 0.05 | 8 | 0.04 | 0.11 | 0.72 |
| Weighted mode | 5 | -0.19 | 0.12 | 0.19 | 3 | -0.14 | 0.15 | 0.43 | 8 | -0.05 | 0.18 | 0.78 |
| Sensitivity test | Estimate |  | p-value/CI |  | Estimate |  | p-value/CI |  | Estimate |  | p-value/CI |  |
| Q (p-value) | 1.65 |  | 0.8 |  | 3.6 |  | 0.17 |  | 23.08 |  | 1.70E-03 |  |
| I2 (CI) | 0.00% |  | (0,0%; 49,5%) |  | 44.30% |  | (0,0%; 83,4%) |  | 69.70% |  | (36,8%; 85,4%) |  |
| Q-Q' (p-value) | 0.46 |  | 0.5 |  | 1.75 |  | 0.19 |  | 1.96 |  | 0.16 |  |
| Egger intercept (p-value) | -0.04 |  | 0.29 |  | -0.01 |  | 0.19 |  | -0.01 |  | 0.75 |  |
| Rucker Test | IVW |  |  |  | IVW |  |  |  | IVW |  |  |  |
| After outlier extraction |  |  |  |  |  |  |  |  |  |  |  |  |
| MR method | Vigorous physical activity → BMI |  |  |  | Moderate physical activity → BMI |  |  |  | Sedentary time → BMI |  |  |  |
|  | SNP | beta | SE | p-value | SNP | beta | SE | p-value | SNP | beta | SE | p-value |
| Egger | NA | NA | NA | NA | NA | NA | NA | NA | 5 | 0.49 | 0.36 | 0.26 |
| Weighted median | NA | NA | NA | NA | NA | NA | NA | NA | 5 | 0.10 | 0.10 | 0.29 |
| IVW | NA | NA | NA | NA | NA | NA | NA | NA | 5 | 0.11 | 0.08 | 0.16 |
| Weighted mode | NA | NA | NA | NA | NA | NA | NA | NA | 5 | 0.08 | 0.14 | 0.59 |
| Sensitivity test | Estimate |  | p-value/CI |  | Estimate |  | p-value/CI |  | Estimate |  | p-value/CI |  |
| Q (p-value) | NA |  | NA |  | NA |  | NA |  | 4.7 |  | 0.32 |  |
| I2 (CI) | NA |  | NA |  | NA |  | NA |  | 15 |  | (0.0%; 82.3%) |  |
| Q-Q' (p-value) | NA |  | NA |  | NA |  | NA |  | 1.41 |  | 0.24 |  |
| Egger intercept (p-value) | NA |  | NA |  | NA |  | NA |  | -1.00E-02 |  | 0.24 |  |
| Rucker Test | NA |  |  |  | NA |  |  |  | IVW |  |  |  |
| BMI, Body mass index (BMI); MR, Mendelian randomization; SNP, single nucleotide polymorphism; SE, standard error; NA, not applicable; IVW, inverse variance weighted; CI, 95% confidence interval. NA: not applicable. Note: No outlier extraction was performed for moderate physical activity → BMI and vigorous physical activity → BMI directions, since Q test and I2 did not indicate heterogeneity. |  |  |  |  |  |  |  |  |  |  |  |  |

**Table S4.** Mendelian randomization results using the IVW, Egger, weighted median and weighted mode methods of BMI on moderate PA, vigorous PA, or sedentary time

|  |  |  |  |  |  |  |  |  |  |  |  |  |
| --- | --- | --- | --- | --- | --- | --- | --- | --- | --- | --- | --- | --- |
| Before outlier extraction |  |  |  |  |  |  |  |  |  |  |  |  |
|  | BMI → Vigorous physical activity |  |  |  | BMI → Moderate physical activity |  |  |  | BMI → Sedentary time |  |  |  |
| MR method | SNP | beta | SE | p-value | SNP | beta | SE | p-value | SNP | beta | SE | p-value |
| Egger | 64 | 0.09 | 0.08 | 0.22 | 65 | 0.17 | 0.08 | 0.04 | 65 | -0.10 | 0.09 | 0.28 |
| Weighted median | 64 | -0.04 | 0.03 | 0.21 | 65 | 0.004 | 0.04 | 0.92 | 65 | 0.03 | 0.04 | 0.44 |
| IVW | 64 | -0.08 | 0.03 | 0.005 | 65 | -0.05 | 0.03 | 0.09 | 65 | 0.07 | 0.03 | 0.02 |
| Weighted mode | 64 | -0.01 | 0.07 | 0.86 | 65 | 0.02 | 0.05 | 0.72 | 65 | -0.05 | 0.10 | 0.59 |
| Sensitivity test |  |  |  |  |  |  |  |  |  |  |  |  |
| Sensitivity test | Estimate |  | p-value/CI |  | Estimate |  | p-value/CI |  | Estimate |  | p-value/CI |  |
| Q (p-value) | 92.4 |  | 9.30E-03 |  | 83.73 |  | 0.05 |  | 97.5 |  | 4.40E-03 |  |
| I2 (CI) | 31.80% |  | 6.9%; 50.00% |  | 23.60% |  | (0.00%; 44.20%) |  | 34.40% |  | 10.80%; 51.70% |  |
| Q-Q' (p-value) | 8.36 |  | 3.80E-03 |  | 10.86 |  | 9.81E-04 |  | 6.59 |  | 1.00E-02 |  |
| Egger intercept (p-value) | -4.99E-03 |  | 9.90E-01 |  | -6.40E-03 |  | 9.81E-04 |  | 4.96E-03 |  | 2.00E-02 |  |
| Rucker test | Egger |  |  |  | Egger |  |  |  | Egger |  |  |  |
| After outlier extraction |  |  |  |  |  |  |  |  |  |  |  |  |
|  | BMI → Vigorous physical activity |  |  |  | BMI → Moderate physical activity |  |  |  | BMI → Sedentary time |  |  |  |
| MR method | SNP | beta | SE | p-value | SNP | beta | SE | p-value | SNP | beta | SE | p-value |
| Egger | 57 | 0.13 | 0.07 | 0.06 | 55 | 0.16 | 0.07 | 0.04 | 57 | -0.02 | 0.08 | 0.76 |
| Weighted median | 57 | -0.04 | 0.03 | 0.27 | 55 | 2.95E-03 | 0.04 | 0.93 | 57 | 0.03 | 0.04 | 0.48 |
| IVW | 57 | -0.07 | 0.02 | 0.01 | 55 | -0.06 | 0.03 | 0.03 | 57 | 0.07 | 0.03 | 0.01 |
| Weighted mode | 57 | -0.01 | 0.06 | 0.83 | 55 | 0.02 | 0.06 | 0.68 | 57 | -0.06 | 0.08 | 0.46 |
| Sensitivity test |  |  |  |  |  |  |  |  |  |  |  |  |
| Sensitivity test | Estimate |  | p-value/CI |  | Estimate |  | p-value/CI |  | Estimate |  | p-value/CI |  |
| Q (p-value) | 61.88 |  | 0.27 |  | 40.51 |  | 0.91 |  | 56.29 |  | 0.46 |  |
| I2 (CI) | 9.50% |  | 0.00% - 35.10% |  | 0.00% |  | (0.0%; 8.9%) |  | 0.50% |  | 0.0%; 31.50% |  |
| Q-Q' (p-value) | 9.79 |  | 1.75E-03 |  | 9.6 |  | 1.94E-03 |  | 1.62 |  | 2.00E-01 |  |
| Egger intercept | -5.70E-03 |  | 1.75E-03 |  | -6.44E-03 |  | 9.15E-05 |  | 2.60E-03 |  | 2.00E-01 |  |
| Rucker test | IVW |  |  |  | IVW |  |  |  | IVW |  |  |  |
| BMI, Body mass index; MR, Mendelian randomization; SNP, single nucleotide polymorphism; SE, standard error; NA, not applicable; IVW, inverse variance weighted; CI, 95% confidence interval. |  |  |  |  |  |  |  |  |  |  |  |  |

**Table S5.**
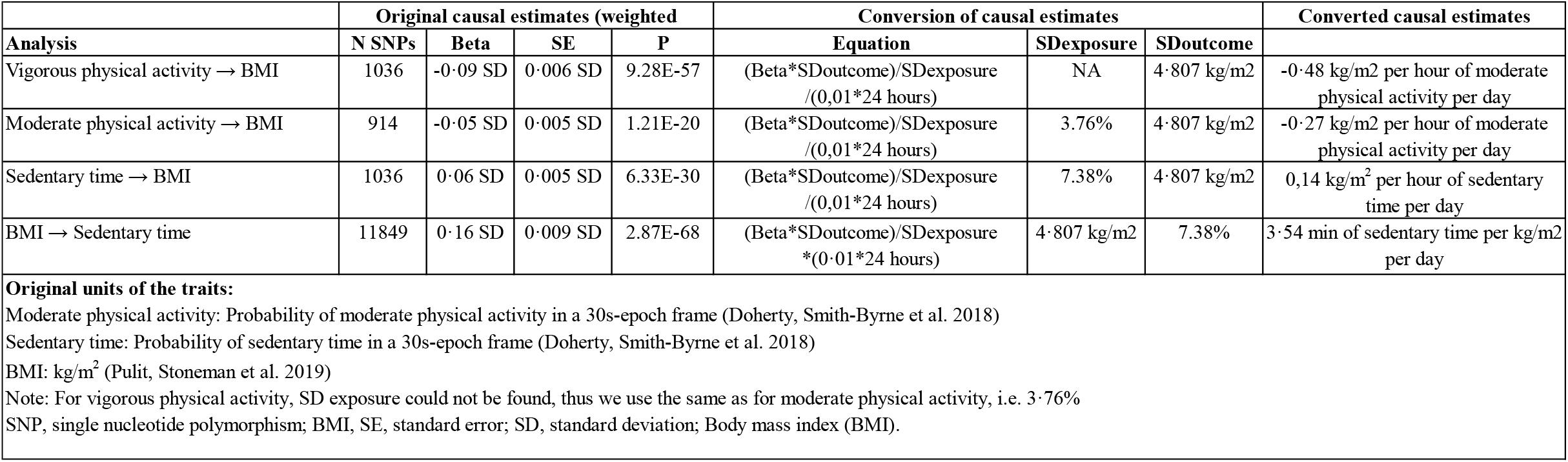
Approximation of the causal estimates in absolute units

## Additional file 2. Analysis plan for causality between physical activity, sedentary behaviour, and obesity: A Mendelian randomization study

Date: initiated in March 2020

Motivation: Obesity is a global epidemic increasing morbidity and mortality worldwide. Physical inactivity and increased sedentary time are associated with excess weight gain as shown from observational studies. However, observational studies suffer from residual confounding and reverse causality. Mendelian randomization helps to overcome confounding and reverse causality by instrumenting the exposure trait using genetic variants. Here, we aim to assess the causality between physical activity, sedentary behaviour and body mass index in adults by bidirectional Mendelian randomization analyses.

Inclusion criteria for genome wide summary level data:

- Largest published genome-wide association study summary statistics
- European ancestry
- Objectively measured continuous traits

Exposures:

- Vigorous physical activity
- Moderate physical activity
- Sedentary time
- Body mass index

Outcomes:

- Vigorous physical activity
- Moderate physical activity
- Sedentary time
- Body mass index

Directions of association for outcome and exposure:

- Vigorous physical activity → body mass index
- Moderate physical activity → body mass index
- Sedentary time → body mass index
- Body mass index → vigorous physical activity
- Body mass index → moderate physical activity
- Body mass index → sedentary time

Mendelian randomization methods:

- The Causal Analysis Using Summary Effect estimates (CAUSE) method
- Inverse variance weighted (IVW)
- Mendelian randomization-Egger (MR-Egger)
- Weighted median
- Weighted mode

Sensitivity tests and plots to account for heterogeneity and horizontal pleiotropy in the IVW, MR-Egger, weighted median and weighted mode analyses:

- Steiger filtering
- Automated outlier removal with RadialMR
- Rucker framework
- Cochran’s Q method
- Leave-one-out forest plots
- Funnel plots

Note: The Mendelian randomization analyses, sensitivity tests, and sensitivity plots for IVW, MR-Egger, weighted median and weighted mode methods were repeated after outlier removal.

Post-hoc analysis:

When less than three genetic variants associated with the exposure trait were available for the IVW, MR-Egger, weighted median, and weighted mode methods using genome-wide significant threshold (P < 5×10-8), we used a p-value threshold of P < 5×10-7 to identify a sufficient number of genetic instruments to produce stable estimates and plots.

R packages used:

CAUSE^1^

TwoSampleMR^2^

RadialMR^3^

Meta R^4^

